# Bicistronic CAR T-cells Against CD70 & Active Integrin β2 Overcome Antigen Heterogeneity and Preserve Safety in Acute Myeloid Leukemia

**DOI:** 10.1101/2025.09.21.677627

**Authors:** Amrik S Kang, Haley Johnson, Nicole Lei, Jeremiah Wong, Nabeel Razi, Adila Izgutdina, Corynn Kasap, Nikhil Chilakapati, Jose Rivera, Paul Phojanakong, Juan Antonio Camara Serrano, Fernando Salangsang, Veronica Steri, Aaron C Logan, Justin Eyquem, Benjamin J Huang, Arun P Wiita

## Abstract

The surface antigen landscape of acute myeloid leukemia (AML) displays significant heterogeneity and overlap with healthy hematopoietic cells. This imparts a substantial hurdle to the development of AML-targeting chimeric antigen receptor (CAR) T-cells that can avoid on-target, off-tumor toxicity. Here, we develop a dual-antigen targeting CAR-T against CD70 and the active conformation of integrin β2 (aITGB2), each previously reported as promising AML targets due to minimal off-tumor expression. We show an OR-gated approach for these antigens significantly increases the proportion of AML blasts that can be targeted, in part using a novel *ex vivo* co-culture method to restore surface protein homeostasis following a freeze-thaw cycle. We test dual-targeting CAR-T constructs with different combinations of costimulatory domains, identifying constructs with superior anti-tumor cytotoxicity *in vitro* against AML cell line and patient-derived xenograft models. We further show significantly improved *in vivo* tumor clearance and survival for a dual-targeting CAR in murine models of AML tumor heterogeneity. Finally, we show that this dual-targeting CAR does not increase off-tumor toxicity, especially against hematopoietic stem and progenitor cells. Together, these findings demonstrate a promising clinically-translatable approach for the treatment of AML without the notable toxicity liabilities associated with other leading CAR-T targets for this disease.

## Introduction

Acute myeloid leukemia (AML) is the most devastating common blood cancer in adults, associated with over 20,000 new diagnoses and 11,000 deaths annually in the United States alone^1^. Despite the notable success of allogeneic hematopoietic stem cell transplant (allo-HSCT) as a potentially curative intervention, many patients are not eligible for transplant or may relapse post-transplant^2,3^. Furthermore, despite the recent commercial availability of several new therapeutic small molecules, these agents have primarily led to modest lifespan extension and not cures^4–6^. As a result, overall 5-year survival rates for AML have remained at approximately 30% for decades^7^. These outcomes underscore the enormous clinical need for new therapies against AML that can induce remission and improve patient survival.

Cellular immunotherapies, including chimeric antigen receptor T-cells (CAR-Ts), have revolutionized the treatment of B-cell leukemia/lymphoma and multiple myeloma^8–12^, but these therapies have not shown success in the clinic for AML^13–21^. While there are many mechanisms that contribute to a lack of CAR T efficacy in AML^22–25^, one widely-acknowledged limitation is “on target, off tumor” toxicity of the leading AML CAR-T targets CD33, CD123, and CLEC12A, which are also expressed on healthy hematopoietic stem and progenitor cells (HSPCs) and/or mature myeloid cells^26–28^. These toxicities may limit the therapeutic dose tolerated in patients and have resulted in severe adverse events and even patient deaths^29–31^. To avoid toxicities, others have proposed complex engineering strategies to tune CAR affinity to avoid healthy tissue antigen levels^32–34^. Alternative approaches have also included pairing immunotherapy treatment with allo-HSCT, sometimes with genetic editing of the target antigen on the transplanted cells to rescue normal hematopoiesis^19,35–37^. While certainly intriguing, this strategy may still not be accessible for transplant-ineligible patients and necessitates two major therapeutic interventions.

Another approach to address this on-target, off-tumor toxicity is identifying alternative AML-specific CAR-T targets. Combined transcriptomic and proteomic analysis from the Sadelain group identified CD70 as a particularly promising target, with notably high expression on both bulk AML tumor cells and leukemic stem cells and an absence of expression on normal bone marrow HSPCs^38^. Alternatively, our group developed the technology “structural surfaceomics”, combining cross-linking mass spectrometry with cell surface proteomics, to identify the active conformation of integrin β2 (aITGB2) as a conformation-selective, AML-specific target that is also not expressed on HSPCs or other mature hematopoietic cells^39^. Other analyses have also identified ADGRE2, CCR1, LILRB2, LILRB4, TIM-3, CD38, IL1RAP, SIGLEC-6, and CD7, among others, as potential AML targets^38,40–46^. However, nearly all of these antigens have only limited expression on AML and/or are also expressed frequently on healthy hematopoietic cells.

Further complicating the issue of immunotherapeutic target selection, AML is well known to feature significant intra-tumoral heterogeneity of surface antigen expression. This dynamic can lead to antigen-negative relapse following single antigen-targeted treatment^38,43,47^. One potential solution is to utilize a multi-antigen targeting strategy, thereby both recognizing a larger percentage of AML blasts and making it more difficult for antigen escape through downregulation of a single antigen. Several analyses have attempted to identify a combination of antigens that would be most suitable for targeting AML without leading to intolerable toxicity, but most combinations still result in targeting of HSPCs or a mature hematopoietic cell lineage^38,43,47^.

Several approaches to dual-targeting CAR-Ts have been evaluated in various tumor models and clinical trials to activate CAR-Ts upon binding to either of two target antigens, commonly referred to as an “OR” logic gate. These include “pooled” CAR-T products consisting of a mixture of T-cells that individually express CARs targeting either antigen of interest, CAR-Ts that express multivalent binders in either a “tandem” or “loop” conformation with a shared transmembrane and signaling domain, and “bicistronic” CAR-Ts that express both CARs on the same cell^48,49^. Although data remains limited, particularly in the context of AML, several studies have shown trends toward the bicistronic approach outperforming other strategies^50,51^. This advantage may be due to more robust CAR signaling from both target antigens compared to tandem CARs and superior protection against antigen-negative relapse compared to the pooled CAR approach^50,51^. Other groups have also studied the feasibility of complex logic-gated targeting of AML target antigens with notable preclinical successes, but this approach generally requires laborious tuning of binder affinities in order to avoid unacceptable off-tumor toxicities, and may not be widely applicable outside of carefully chosen AML models^32,43,52,53^.

Here, we design an OR logic-gated bicistronic CAR to target two antigens, CD70 and aITGB2, which are thought to be largely absent from HSPCs and normal tissues. This approach minimizes off-tumor toxicity while significantly increasing AML blast targeting when compared to single-target CAR-Ts. We show that a bicistronic CAR design that incorporates both a CD28 and 4-1BB costimulatory domain (one on each individual CAR) shows superior performance in several AML models, both *in vitro* and *in vivo*, utilizing both cell lines and patient-derived xenografts (PDXs). Furthermore, we show that this bicistronic CAR design maintains no appreciable toxicity against HSPCs, similar to its individual component CARs, demonstrating its promise as a clinically translatable therapy for AML that can avoid major toxicities seen in other AML-targeting CAR-Ts.

## Results

### CD70 and active ITGB2 are a logical pairing for AML targeting

To validate the selection of the cell surface antigens CD70 and aITGB2, we first used mRNA sequencing data to look at the transcript levels for both antigens in three AML cohorts: Beat AML (adult)^54^, the Cancer Genome Atlas (TCGA) (adult)^55^, and TARGET AML (pediatric)^56^. In all three analyses, significant, albeit heterogeneous, transcript levels were seen for both targets on AML blasts (**Fig. 1A**), with the caveat that an RNA-level analysis could only measure transcript levels for total *ITGB2*, rather than the AML-specific active conformation. We then compared the expression of these two antigens, as well as the well-studied AML targets *CD33, IL3RA* (CD123), and *CLECL12A* (CLL-1), in AML blasts to their expression in normal CD34-positive cells from bone marrow. While *CD33, IL3RA, and CLEC12A* each had notably positive expression in the healthy CD34+ HSPCs, we observed nearly no expression of *CD70* in normal tissue in all three cohorts (**Fig. 1B**), highlighting its lower potential for off-tumor toxicity. As expected, *ITGB2* was seen at high levels in both tumor and healthy HSPCs due to the inability to differentiate the active and inactive forms of the target antigen. We further compared levels of *CD70* and *ITGB2* expression from patient bulk bone marrow samples in the TARGET cohort collected at initial diagnosis compared to matched samples at the end of induction chemotherapy. We observed a significant decrease in *CD70* at the second time point, consistent with a reduction in tumor burden. This decreased expression was not seen in transcripts encoding *CD33* and *IL3RA* (**Fig. 1C**). This result suggests that while the majority of *CD70* expression could be attributed to tumor cells, most *CD33*- and *IL3RA*-expressing cells were likely healthy tissue unaffected by chemotherapy. Together with the results in **Fig. 1B**, this analysis supports the potential for a CD70-targeting therapy to have a wider therapeutic index to avoid the on-target, off-tumor toxicities seen in CD33-, IL3RA-, or CLEC12A- targeting CAR-T therapies.

**Figure 1.**
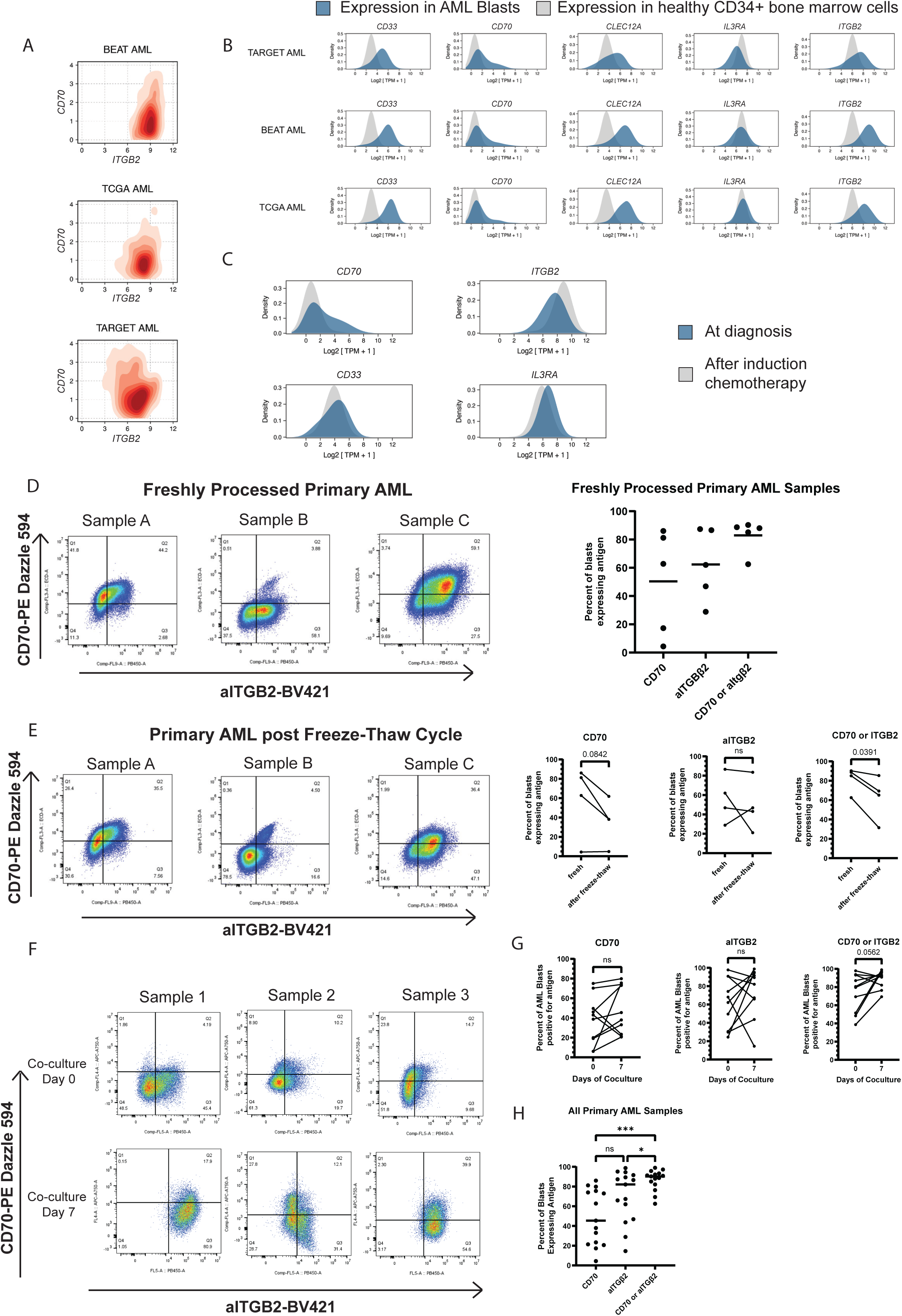
CD70 and aITGB2 are a good antigen pair for OR-gated AML therapeutics: **A)** Bulk RNA-seq analysis from three different AML patient sample cohorts for gene co-expression of *CD70* and *ITGB2*, with darker orange coloring representing a greater density of patient samples. **B)** Bulk RNA-seq analysis for relative expression of major AML targets in AML blasts (blue) and healthy CD34+ bone marrow (grey) in three AML patient cohorts. **C)** Bulk RNA-seq analysis for the relative expression of AML targets in patient bone marrow samples obtained at the time of diagnosis with AML (blue) and after the completion of induction chemotherapy (grey). **D)** *Left:* Representative flow cytometry plots and gating for CD70 and aITGB2 surface expression on freshly processed AML samples; *Right:* Percent of AML blasts expressing CD70 and/or aITGB2 on all N = 5 freshly processed AML samples. **E)** *Left:* Flow cytometry plots of matched samples from Fig 1D immediately following one freeze-thaw cycle; *Right:* Change in percent of AML blasts expressing CD70 and/or aITGB2 in matched samples following a freeze-thaw cycle, 2-sided paired T-test. **F)** Representative flow cytometry plots of frozen AML primary samples immediately after initial thaw (top) and following 7 days of co-culture with HS-5 bone marrow stromal cell line and supportive cytokines (bottom). **G)** Changes in antigen expression for each of N = 10 frozen AML samples following 7 days of stromal cell line and supportive cytokine co-culture, 2-sided paired T-test. **H)** Distribution of antigen expression in N = 15 AML combined fresh and co-cultured frozen primary samples, Tukey’s multiple comparisons test, one-way ANOVA test; * = p < 0.05, *** = p < 0.001

To profile the status of aITGB2 on AML and determine its co-occurrence with CD70 on individual blasts, we next performed flow cytometry analysis on a limited number of fresh AML patient samples. We observed a similar pattern to the RNA-seq analysis of significant but heterogeneous expression of these antigens, with improved tumor coverage when targeting CD70 or aITGB2 (**Fig. 1D**, **Supp. Fig. 1A**). To expand our sample size, we next attempted to profile banked, frozen AML samples, but quickly noted that expression levels of both antigens appeared to artifactually decrease immediately following a freeze-thaw cycle (**Fig. 1E**), potentially due to shedding or internalization of surface proteins during the thaw process.

Therefore, we developed an *ex vivo* culture system for frozen primary AML samples. Our goal was to better simulate the *in vivo* bone marrow microenvironment, both to promote AML tumor growth and re-establish cell surface protein homeostasis. Primary samples were thawed and maintained in RPMI-1640 media with added 20% (v/v) FBS, 25 ng/mL interleukin-3 (IL-3), stem cell factor (SCF), and Fms-related tyrosine kinase 3 ligand (FLT3L), and co-cultured for up to 7 days with the human bone marrow stromal cell line HS-5 (**Supp. Fig. 1B**). By flow cytometry, we observed a moderate increase in CD70 and aITGB2 expression in a subset of primary samples during the co-culture period (**Fig. 1F-G**, **Supp. Fig. 1C**). We specifically observed that the expression of CD70 and/or aITGB2 following the stromal cell co-culture system closely matched that of the fresh (not previously frozen) primary AML samples previously analyzed (**Supp. Fig. 1D**). Therefore, we propose that our *ex vivo* system was able to largely restore physiological surface expression of our proteins of interest on blasts after freeze-thaw.

With this expression data in hand, we then evaluated whether a dual-targeting approach for these two antigens might improve the ability for a targeted therapy to recognize a greater proportion of AML blasts in these primary samples when compared to targeting either antigen on its own. We observed that, when considering fresh and previously frozen AML primary samples together, a median 90.3% (IQR 82 - 93%) of AML blasts could be targeted using an OR-gated targeting approach for CD70 and aITGB2, significantly greater (p= 0.0067, RM one-way ANOVA) than could be achieved by targeting aITGB2 alone (median 82.1%, IQR 47 - 92%) or CD70 alone (median 45.5%, IQR 21-76%) (**Fig. 1H).** These data illustrate the potential for an OR-gated CAR-T against both antigens to lead to significantly improved AML tumor control when compared to single antigen-targeting CAR Ts.

### Dual-targeting CAR construct design and validation

Given these expression patterns, we next sought to design a dual-targeting, OR-gated CAR T against aITGB2 and CD70. As both surface antigens are known to be moderately upregulated on activated T-cells^39,57,58^, we first optimized a CRISPR Cas9-based gene editing approach that would prevent cross-activation and potential fratricide by CAR-expressing cells during the manufacturing process. We therefore evaluated a set of 2 *CD70-* and 4 *ITGB2*-targeting guide RNAs (gRNAs) on CD3-isolated donor T-cells for their ability to genetically knock out both targets. We achieved >90% KO for *CD70* and >80% KO for *ITGB2* with our best-performing gRNAs without the need for enrichment (**Supp. Fig. 2A**). We observed no defect in T-cell proliferation in edited cells (**Supp. Fig. 2B**), in agreement with previous data on editing these genes individually in T-cells ^39,59^.

We next generated and tested multiple designs for a dual CD70 & aITGB2-targeting CAR. We used a bicistronic format with two independent second-generation CARs encoded on the same plasmid. As noted above, this design has previously been shown to outperform other dual-targeting CAR formats^50,51^ and ensures independent activation of the CAR T-cell by the presence of either antigen. We tested eight bicistronic CAR designs (denoted as bicistronic CAR design (BCD) 1-8) utilizing our previously characterized binders against aITGB2^39^ and CD70^59^, differing in the order on the plasmid in which the CARs are expressed and in the combination of costimulatory domains (CD28- or 4-1BB-based) expressed by each CAR (**Fig. 2A**). We observed moderate differences in the transduction efficiency of the different designs, as measured by the co-expression of extracellular myc- and FLAG- tags on each expressed CAR, despite their shared binder sequence identity (**Fig. 2B**). The highest transduction efficiency was observed in construct BCD 1, 5, and 6. As has been previously reported^60–62^, the CAR construct that was placed second in the bicistronic construct, after a T2A ribosomal skipping sequence, was observed to have moderately lower overall expression compared to the construct that was placed before the T2A, regardless of the CAR identity.

**Figure 2.**
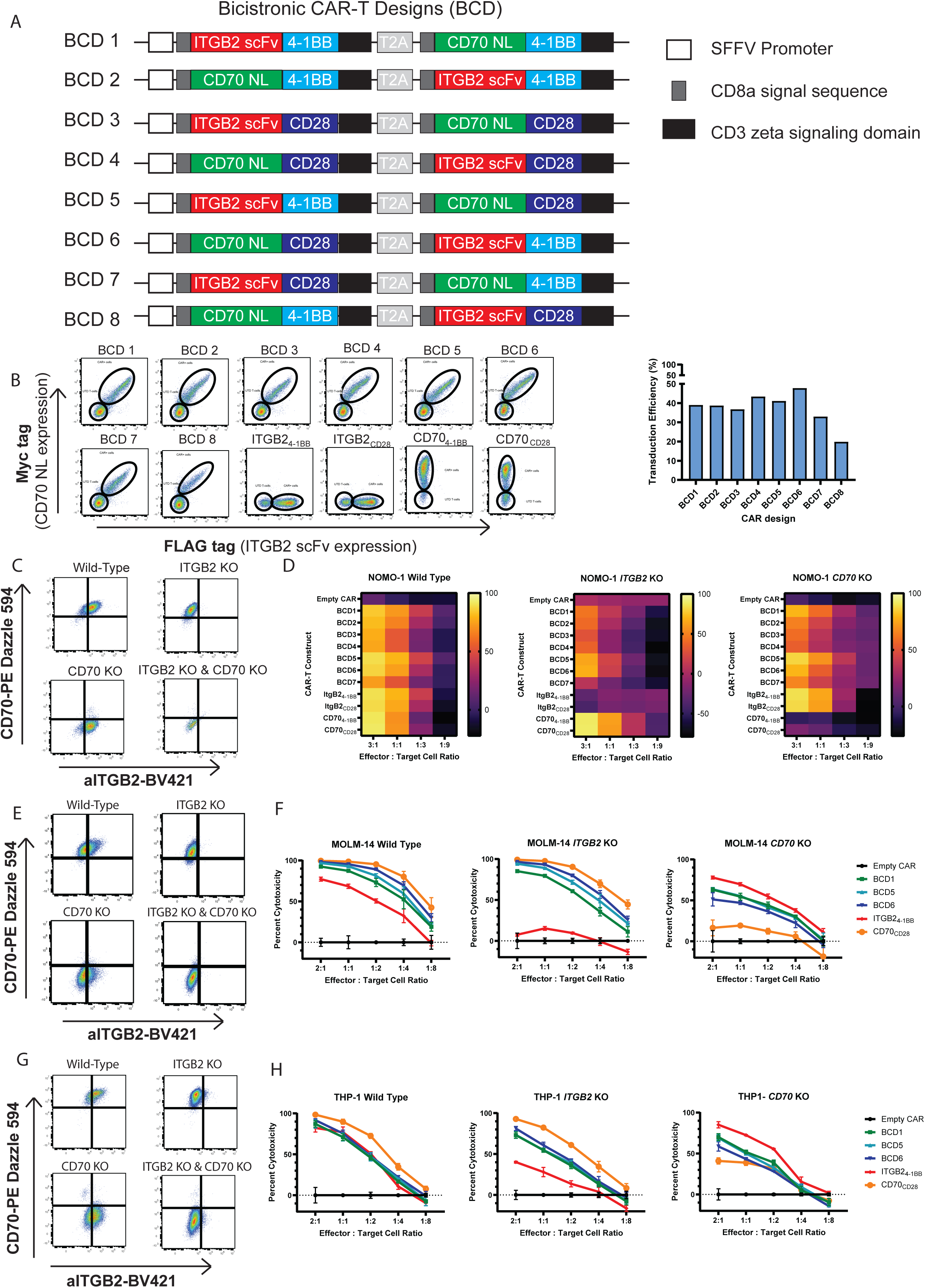
Bicistronic CAR-Ts targeting CD70 & aITGB2 effectively eliminate AML cell lines *in vitro*: **A)** Cartoon depiction of all bicistronic dual-targeting CAR-T constructs that were cloned and tested for *in vitro* cytotoxicity. **B)** *Left:* Representative flow cytometry plots of co-expression of each CAR construct; *Right:* Transduction efficiency of all bicistronic CAR constructs tested. **C)** Expression of CD70 and aITGB2 in the wild-type, ITGB2 knockout, CD70 knockout, and dual-antigen knockout cell lines derived from NOMO-1. **D)** Heat map of relative antitumor cytotoxicity in a 24 hour luciferase cytotoxicity assay in WT and KO NOMO-1 lines. CAR constructs were tested at 4 effector-to-target (E:T) ratios and displayed as averages of N = 3 replicates for each condition. Number of T-cells added for each concentration were normalized for transduction efficiency. **E)** Expression of CD70 and aITGB2 in the wild-type, ITGB2 knockout, CD70 knockout, and dual-antigen knockout cell lines derived from MOLM-14. **F)** Relative antitumor cytotoxicity in WT and KO MOLM-14 lines by CAR constructs in a 24 hour luciferase assay. Each CAR was tested at 5 E:T ratios and displayed as averages of N = 3 replicates for each condition, error bars represent SEM. **G)** Expression of CD70 and aITGB2 in the wild-type, ITGB2 knockout, CD70 knockout, and dual-antigen knockout cell lines derived from THP-1. **H)** Relative antitumor cytotoxicity in WT and KO THP-1 lines by CAR constructs in a 24 hour luciferase assay. Each CAR was tested at 5 E:T ratios and displayed as averages of N = 3 replicates for each condition, error bars represent SEM.

We then tested these eight bicistronic CAR designs head-to-head for their ability to kill AML tumors at multiple effector-to-target cell (E:T) ratios using cell line models that express both target antigens. We performed our initial screen using the NOMO-1 cell line due to its strong expression of both CD70 and aITGB2. To ensure that both CARs in each construct were capable of driving specific tumor killing, we first generated genetic knockouts of *CD70* or *ITGB2* for NOMO-1 and performed FACS sorting to ensure pure antigen KO populations (**Fig. 2C**). We then performed 24-hour cytotoxicity assays with the cell line and its KO variants by co-culturing them with either a bicistronic CAR variant, a non-targeting CAR lacking an antigen recognition domain (“empty CAR”), or single-antigen targeting CARs against CD70 alone or aITGB2 alone. We consistently observed that best performing dual-antigen targeting CARs were BCD 1, 5 and 6 (**Fig. 2D**), which were then selected for additional screening and characterization. We next performed the same KO and co-culture cytotoxicity experiments with the AML cell lines MOLM- 14 and THP-1 (**Fig. 2E-H; Supp. Fig . 2C**), observing that each dual-targeting CAR maintained effective, dose-dependent killing even after one of the two targeted antigens had been genetically knocked out, while the single-antigen targeting CARs showed essentially no cytotoxicity against their respective KO cell line, highlighting the independent efficacy of both CARs.

### Dual-targeting CARs display potent antitumor efficacy *in vivo* in a model of AML antigen heterogeneity

Following the identification of our most promising bicistronic CAR constructs, we next wanted to test their ability to control AML tumors in an *in vivo* context. To ensure we tested the activity of both CAR constructs present in the bicistronic CARs, we opted to utilize a cell line-based tumor heterogeneity model. We intravenously injected NSG mice with 1 million luciferase-expressing THP-1 cells as a 50:50 ratio of *ITGB2* KO cells to *CD70* KO cells to model the antigen heterogeneity of a naturally occurring AML tumor. This design ensured that relapse of antigen-negative tumor would occur unless the CAR-Ts being tested were able to target both antigens for tumor control. Three days following tumor injection, we infused 1.5 million CAR-positive T-cells in each mouse and performed weekly bioluminescence imaging (BLI) and weekly or biweekly peripheral blood draws to monitor CAR proliferation (**Fig. 3A**).

**Figure 3.**
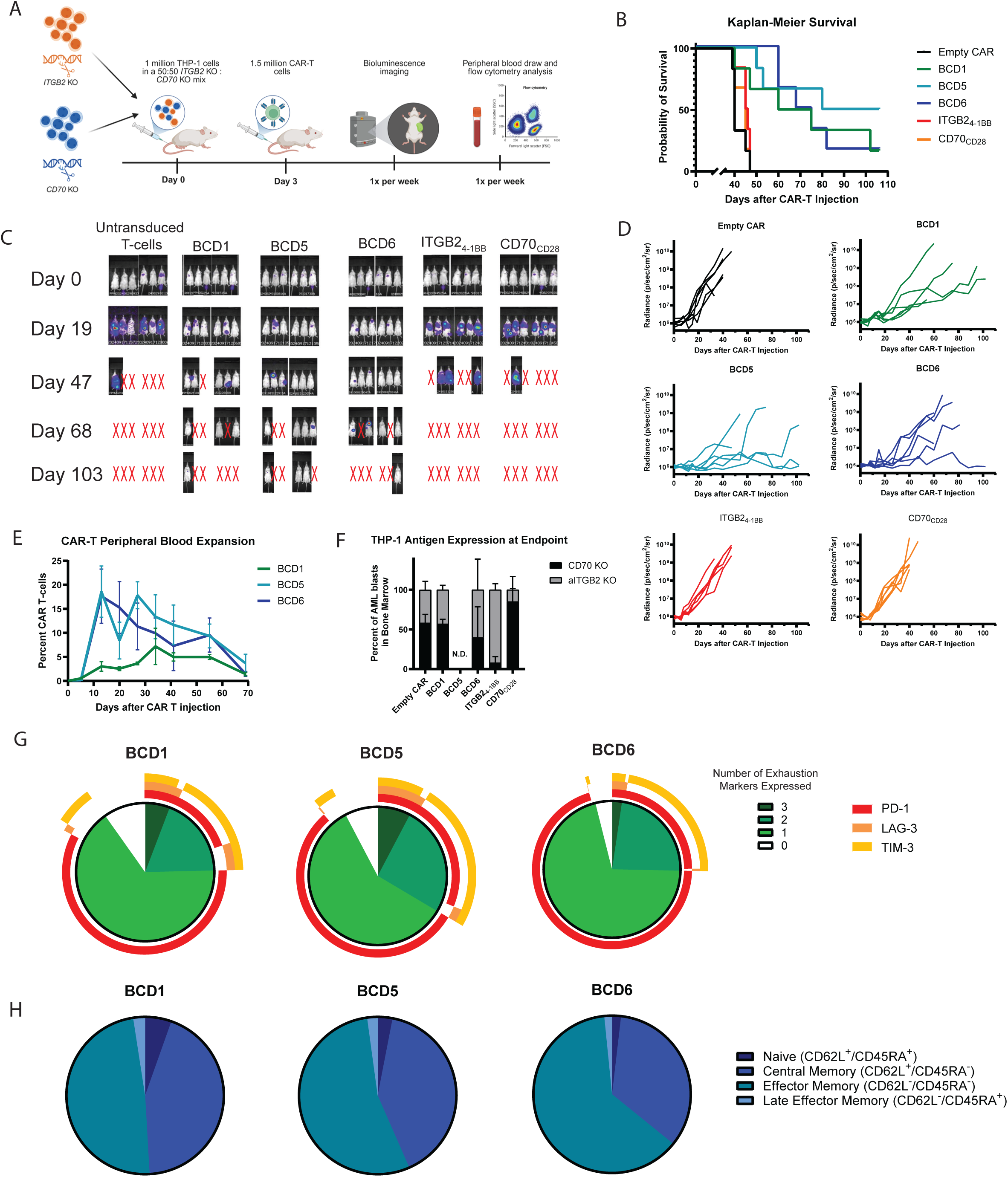
Bicistronic CAR-Ts show superior tumor control in an *in vivo* model of AML antigen heterogeneity: **A)** Timeline of AML antigen heterogeneity xenograft experiment, N = 6 mice per treatment group. **B)** Kaplan-Meier Survival curve of mice by treatment group**. C)** Bioluminescence imaging of mice in each treatment group at several time points following IV injection of d-luciferin. Red Xs indicate an individual mouse that has died or reached a humane endpoint. **D)** Quantification of bioluminescence signal in individual mice over time. **E)** Proliferation of CAR-Ts in each treatment group from weekly peripheral blood draws, as measured by flow cytometry, as a percent of all live singlet-gated events. N = 3 mice evaluated at each time point; error bars represent standard deviation. **F)** Relative proportion of each KO THP-1 line found in the bone marrow at clinical endpoint, as assessed in N = 3 mice per group by flow cytometry, error bars represent standard deviation. **G)** Relative proportion of peripheral blood CAR-T cells expressing the T-cell exhaustion markers PD-1, TIM-3, and LAG-3, 20 days after CAR-T infusion into mice. External rings represent the proportion of CAR-Ts expressing each individual marker within each pie chart subgroup. Data presented is the average of N = 3 mice evaluated per treatment group. **H)** Relative proportion of CAR-Ts classified into each memory/effector T-cell phenotype, 20 days after CAR-T infusion into mice. Classification assessed by flow cytometry of CD62L and CD45RA expression on CAR-T cells isolated from peripheral blood. Data presented is the average of N = 3 mice evaluated per treatment group.

Encouragingly, and consistent with our earlier results, we observed highly effective tumor control in each of our three bicistronic CARs tested, while our negative control “empty” CAR and single antigen-targeting anti-CD70 and anti-aITGB2 CAR-treated mice all died of tumor burden between days 40 and 50 of the study. In comparison, our best-performing bicistronic construct, BCD5, displayed a median survival of 93 days, with 3 of 6 mice still alive at the conclusion of the study (**Fig. 3B-D**). We also profiled antigen expression on isolated tumor from mouse bone marrow at terminal endpoints, observing a skew toward antigen-negative relapse for each of the single antigen-targeted CARs, while the empty CAR and bicistronic CAR conditions showed a mixed population of tumor antigen expression, reflecting expected tumor selection trends (**Fig. 3E**).

Notably, CAR-T expansion seemed correlated with prolonged survival, with BCD5 also being measured at the highest relative levels in the peripheral blood at most time points, peaking at 2 weeks after injection as a mean 18.6% of all non-erythrocyte blood cells (**Fig. 3F, Supp. Fig. 3A**). Interestingly, BCD1, which contains the 4-1BB costimulatory domain on both CARs, appeared to have both a slower initial onset of CAR expansion, taking 4 weeks to reach maximal expansion, and a lesser absolute CAR-T number, which may have contributed to its inferior tumor control in this experiment. The majority of CAR T-cells observed were CD4^+^, with no significant differences in CD4/CD8 ratios observed between the dual-targeting CAR constructs (**Supp. Fig. 3B**). We additionally measured the expression of canonical T-cell exhaustion markers on peripheral blood CAR-Ts at week 3 of the experiment, shortly after the time of peak expansion, observing similar levels of PD-1, Lag-3, and Tim-3 expression between the three bicistronic constructs (**Fig. 3G**). We also explored memory and effector phenotype markers on these CAR-Ts, observing a predominantly central memory (CD62L+/CD45RA-) or effector memory (CD62L-/CD45RA-) phenotype throughout the length of the study, with a shift to the latter over time, with no significant difference observed between the bicistronic constructs (**Fig. 3H, Supp. Fig. 3C**).

### Dual-targeting CARs outperform single antigen-targeting CARs in an AML PDX model

As an orthogonal approach to assess CAR-T effectiveness against AML heterogeneity, we next chose to compare our best-performing dual-targeting CAR, BCD5, against single antigen-targeting CARs in patient-derived xenograft (PDX) models of AML. We first profiled the expression of surface target antigens in two PDXs (referred to here as PDX A and PDX B, used in our prior study^39^ and initially obtained from PRoXe^63^) freshly harvested from previously-implanted mouse bone marrow and spleen. We observed that both PDXs expressed detectable levels of CD33 on nearly all blasts (**Fig. 4A**). We also noted that while PDX B appeared to be fully targetable utilizing a dual-antigen targeting approach, only about 50% of PDX A expressed either CD70 or aITGB2 (**Fig. 4B**). We then repeated our *ex vivo* co-culture experiment using these PDX samples as in **Fig. 1E**, noting that a freeze/thaw cycle led to a moderate decrease in the cell surface levels of both CD70 and aITGB2, which were thereby rescued over a multiday co-culture with HS-5 stromal cells and supportive cytokines and media (**Fig. 4C-D**).

**Figure 4.**
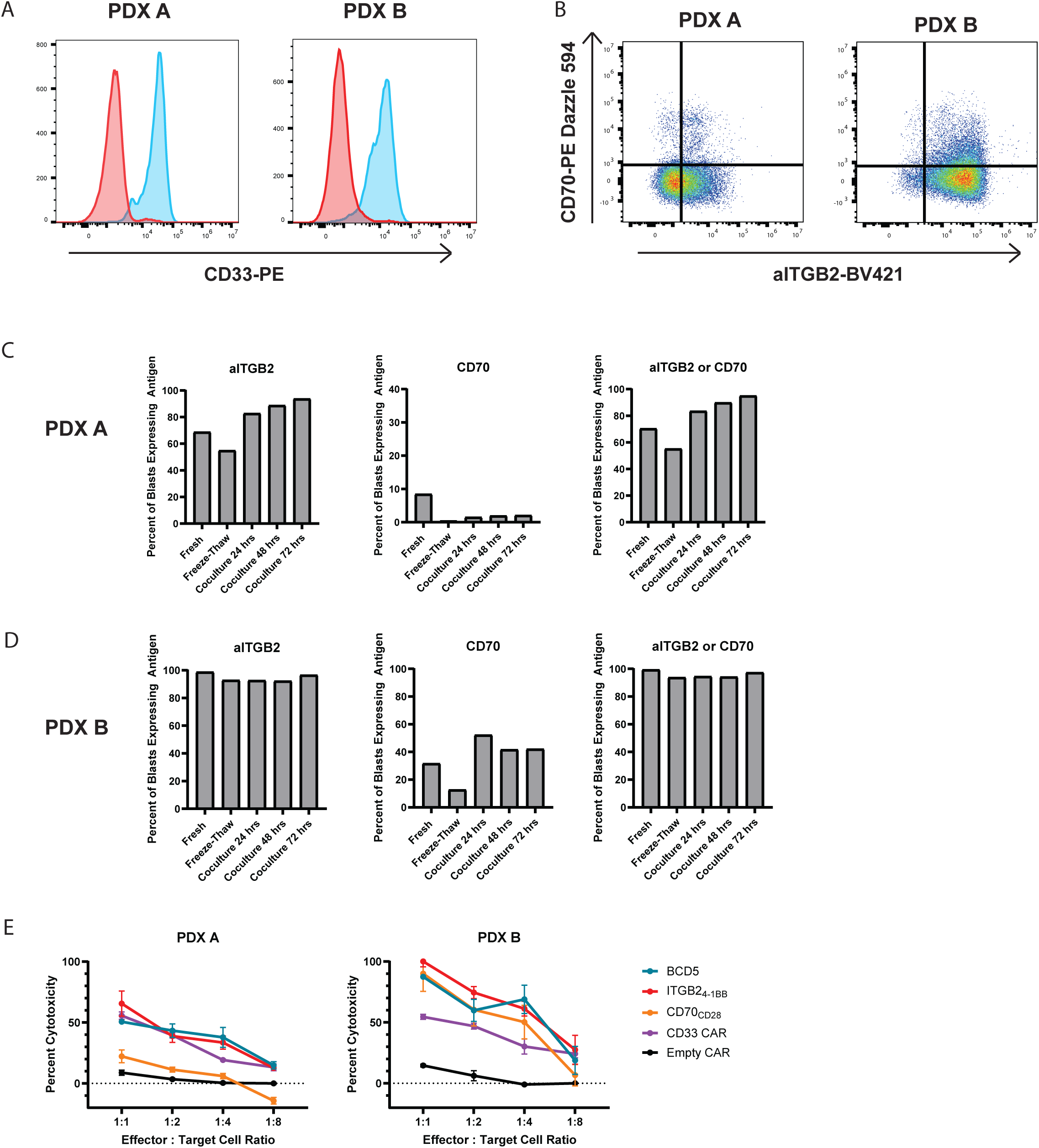
BCD5 effectively targets AML patient-derived xenografts: **A)** Flow cytometry plots of CD33 surface antigen expression in two AML patient-derived xenografts, denoted PDX A and B. **B)** Flow cytometry plots of aITGB2 and CD70 expression in PDX A and B. **C-D)** Expression of CD70 and/or aITGB2 in PDX A **(Fig. C)** and B **(Fig. D)** immediately after thaw, following a freeze-thaw cycle, and after a coculture with bone marrow stromal cell line and supportive cytokines, as previously seen in Fig. 1 D-G. **E)** Relative antitumor cytotoxicity against PDX A and B by CAR constructs in a 24-hour fluorescence-based co-culture assay, tested at 4 E:T ratios and displayed as averages of N = 3 replicates for each condition; error bars represent SEM.

We next tested the *in vitro* antitumor cytotoxicity of BCD5 against both PDX models, comparing it to a negative control empty CAR and single antigen-targeting CARs against CD70, aITGB2, and CD33 (based on the My96 clone binder^64^). As expected based on the antigen expression data, PDX A displayed a maximum of 50-60% cytotoxicity at our highest E:T ratio (1:1) for BCD5, anti-aITGB2, and anti-CD33 CARs, but nearly no cytotoxicity when cocultured with anti- CD70 CARs. In comparison, we saw nearly complete killing of PDX B at the highest (1:1) E:T ratio for BCD5, anti-CD70, and anti-ITGB2 CARs, even outperforming the anti-CD33 CAR (**Fig. 4E**).

Based on the antigen profiling and *in vitro* data, we decided to move forward with PDX B for testing BCD5’s efficacy against an AML PDX *in vivo.* Following busulfan preconditioning, we injected 2 million PDX cells IV into NOG-EXL mice^65,66^, followed by injection of 1.5 million CAR-T cells 10 days later (**Fig. 5A**). We monitored tumor burden and CAR-T expansion through weekly peripheral blood draws until a humane endpoint was reached. In this aggressive tumor model, we observed that BCD5 provided a significant survival benefit compared to empty CAR and anti-CD70 CAR, and very similar survival compared to anti-aITGB2 CAR (**Fig. 5B**). This correlated well with a significantly reduced observable tumor burden in the peripheral blood compared to the empty CAR and anti-CD70 single CAR conditions, with similar performance to the anti-aITGB2 single CAR (**Fig. 5C**), consistent with the antigen expression patterns and *in vitro* activity data.

**Figure 5.**
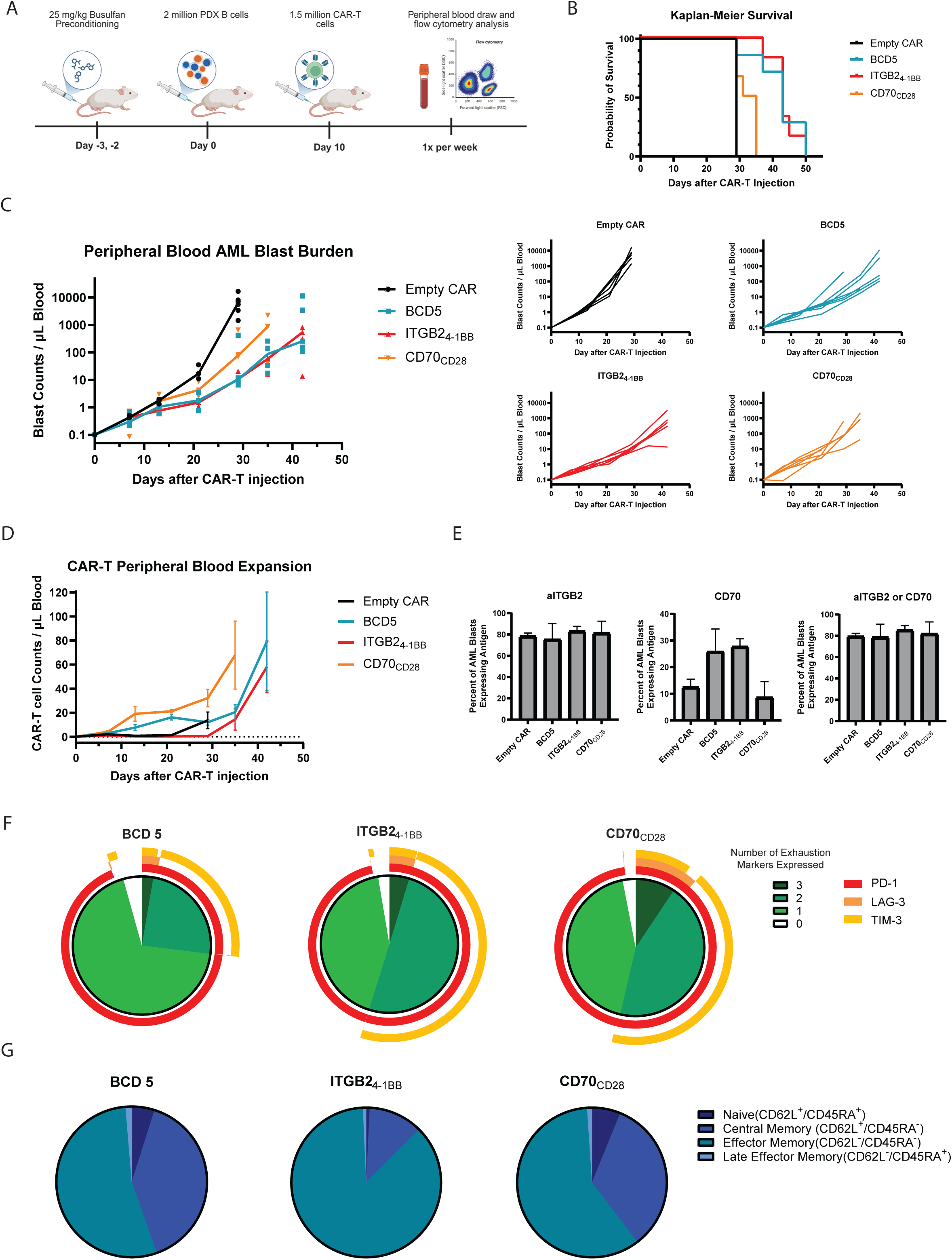
BCD5 shows significant antitumor effect and superior T-cell phenotypes in an *in vivo* AML PDX model: **A)** Timeline of AML PDX xenograft experiment, N = 6 mice per treatment group. **B)** Kaplan-Meier Survival curve of mice by treatment group**. C)** Quantification of AML blast burden, as evaluated by flow cytometry on weekly peripheral blood draws. *Left:* Average blast burden for each group. *Right:* Blast burden for individual mice in each treatment group. **D)** Proliferation of CAR-Ts in each treatment group, as measured by flow cytometry, as a percent of all live singlet-gated events. N = 3 mice evaluated at each time point; error bars represent standard deviation. **E)** Relative proportion of PDX blasts expressing aITGB2 and/or CD70 in the peripheral blood, as assessed in N = 3 mice per group. Data collected by flow cytometry on Day 29 after initial CAR-T treatment; error bars represent standard deviation. **F)** Relative proportion of peripheral blood CAR-T cells expressing the T-cell exhaustion markers PD-1, TIM-3, and LAG-3, 35 days after CAR-T infusion into mice. External rings represent the proportion of CAR-Ts expressing each individual marker within each pie chart subgroup. Data presented is the average of N = 3 mice evaluated per treatment group. **G)** Relative proportion of CAR-Ts classified into each memory/effector T-cell phenotype, 35 days after CAR-T infusion into mice. Classification assessed by flow cytometry of CD62L and CD45RA expression on CAR-T cells isolated from peripheral blood. Data presented is the average of N = 3 mice evaluated per treatment group.

Strikingly, we noted the BCD5 CAR displayed notably more potent expansion in the peripheral blood compared to anti-aITGB2 CAR, with a median 14.9 CAR-Ts/μL detected in the peripheral blood 3 weeks after CAR-T infusion compared to only 0.3 CAR-Ts/μL for the anti-aITGB2 CAR, and a higher peak CAR detection at week 5 as well (**Fig. 5D**). This expansion pattern was notably more similar to the anti-CD70 natural ligand CAR, which has been previously noted to display exceptionally strong *in vivo* expansion^59,67,68^. Interestingly, this similarity also extended to the CD4/CD8 ratio over time, which was CD4^+^ T-cell dominant at later time points for both the BCD 5 and CD70_CD28_ constructs, but which remained CD8^+^ dominant for the ITGB2_4-1BB_ construct (**Supp. Fig. 4A**)

We harvested mouse spleens at the time of terminal sacrifice to characterize antigen expression on detectable PDX blasts. We observed a moderate decrease in CD70 expression in the anti- CD70 CAR condition, but no significant downregulation of aITGB2 in either the anti-aITGB2 CAR or BCD5 mice (**Fig. 5E**). This antigen-positive relapse suggests the mechanism by which the CARs failed to control the tumor was most likely related to impaired CAR-T cell function or an exceptional rate of tumor growth that surpassed the CAR-Ts’ killing capacity, rather than antigen loss or downregulation.

Finally, we noted a less terminally exhausted phenotype of the BCD5 CAR, as measured by expression of PD-1, TIM-3, and LAG-3, when compared to the anti-aITGB2 and anti-CD70 CAR, primarily driven by lower TIM-3 expression for BCD5 (**Fig. 5F**). We also saw a less terminally differentiated phenotype, as measured by expression of CD62L and CD45RA, compared to the anti-aITGB2 CAR at week 5, with a relatively similar split between central and effector memory phenotypes in BCD5 CAR-Ts (**Fig. 5G**). These characteristics are generally thought to be associated with improved CAR-T function and persistence, consistent with the observed expansion advantage. Although these advantages did not translate to improved survival compared to anti-aITGB2 CAR in this model, this is likely an artifact of the PDX B model, which tends to relapse very aggressively at later time points, potentially overcoming some of the inherent differences in tumor killing ability between CAR constructs.

### Dual-targeting CARs display no on-target, off-tumor toxicity against HSPCs

One of the previously noted challenges with developing CAR-Ts against highly expressed AML target antigens is the high likelihood of on-target, off-tumor toxicity against healthy hematopoietic cells, especially hematopoietic stem and progenitor cell (HSPC) populations.

Notably, anti-CD33 CARs, the most widely developed anti-AML CAR-T therapies, have been noted to cause severe depletion of HSPC populations, sometimes leading to severe cytopenias in patients^27,64,69^. To test whether our dual-targeting construct could avoid these toxicities, we first isolated CD34+ HSPCs from a healthy donor by GM-CSF mobilization, then performed a 24-hour co-culture with our BCD5 CAR, single antigen-targeted CARs against CD70, aITGB2, and CD33, and a negative control empty CAR. As expected, we observed a significant reduction in CD34+ cells in the CD33 CAR condition, but no significant effect for either the BCD5 CAR or its constituent single CARs compared to the empty CAR (**Fig. 6A**).

**Figure 6.**
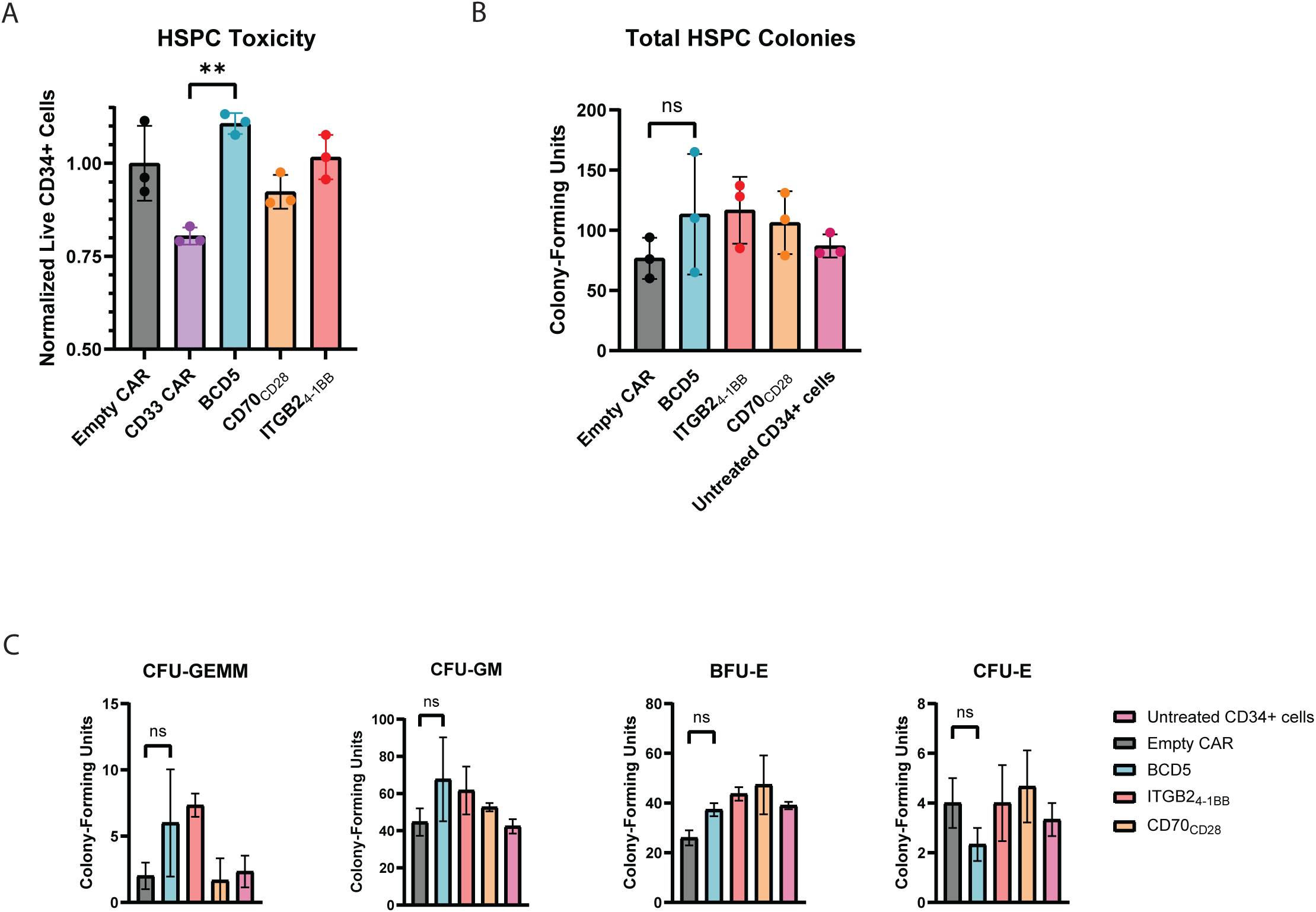
BCD5 shows no signs of toxicity against hematopoietic stem and progenitor cells: A) Relative percent viability of CD34+ HSPCs isolated from healthy GM-CSF mobilized blood calculated via flow cytometry after 24-hour co-culture with CAR constructs. CAR-Ts were cultured at a 3:1 E:T ratio and are displayed as averages of N = 3 replicates for each condition; error bars represent SEM. Tukey’s multiple comparisons test, one-way ANOVA test; ** = p < 0.01. B) Total number of HSPC-derived colonies counted following co-culture with CAR-T constructs and 14 days of colony formation in methylcellulose media. N = 3 replicates per treatment group, error bars represent SEM. Tukey’s multiple comparisons test, one-way ANOVA test. C) Breakdown of HSPC-derived colony subtypes counted following CAR-T coculture and methylcellulose colony formation. N = 3 replicates per treatment group, error bars represent SEM. Tukey’s multiple comparisons test, one-way ANOVA test.

We next performed a long-term CD34+ cell colony-forming assay with the BCD5 CAR and single antigen-targeting CARs to look for any potential toxicity against hematopoietic progenitor cells during early differentiation. Following a 15-day co-culture and colony differentiation process, no impacts were seen on total colonies or colony subsets by the BCD5 CAR or its associated single-targeting CARs when compared to a non-targeting empty CAR or CD34+ cells alone (**Fig. 6B**), highlighting the translational potential for this CAR construct to avoid the HSPC toxicity seen with most other anti-AML CAR-T designs.

## Discussion

Our work presented here demonstrates a novel combination of target antigens for acute myeloid leukemia. AML has had repeated clinical trial failures of potential CAR-T therapies^13–21^. While this lack of clinical success may due to several reasons^22–25^, a widely-acknowledged challenge is both heterogeneity of target antigen expression on AML blasts^38,43,47^ as well as high levels of leading target antigens on healthy tissues, especially HSPCs, resulting in significant risk of “on-target, off-tumor” toxicities^26,27,29^. Here, by prioritizing reduced off-tumor toxicity instead of tumor antigen coverage for each individual CAR, we generated a therapy that targets CD70 and aITGB2 with exceptionally high antigen coverage for AML blasts and no observed toxicity against HSPC populations.

As one of our targets, aITGB2, is a conformation-specific antigen present on AML blasts, its quality as an AML target cannot be determined using traditional RNA-sequencing analyses of large patient cohorts. Instead, profiling this target requires surface-protein level data, such as flow cytometry, which we show here can cause aberrant results if frozen AML patient samples are thawed and directly assayed for expression. Instead, we develop a novel co-culture method with a bone marrow stromal cell line to first allow for an *ex vivo* return to surface expression homeostasis for both antigens, which then allows for an antigen expression profile that more closely resembles the results seen in freshly processed patient AML samples. The expression levels of aITGB2 we report here on primary AML samples also closely align with our previously reported data on the expression of this target antigen^39^. This method can also be explored in the future to evaluate the surface expression of other AML target antigens that are less abundant on the surface, especially if protein-level data is essential to capture target specificity, as is the case here.

CD70 has also been previously identified as a potential target for single-targeted therapies against AML, including for CAR-T and monoclonal antibody therapeutics^67,68,70,71^. Although earlier transcriptomic and proteomic analyses suggested that CD70 could be detected on the surface of nearly all AML blasts by flow cytometry in primary patient samples, most notably by Perna and colleagues in 2017^38^, our data suggests CD70 is detectable, on average, on under 50% of AML blasts, which is more in line with more recent analyses of CD70 expression on AML primary samples^68,72,73^. This finding further supports utilizing a combined targeting approach, such as the bicistronic CAR strategy used here, rather than monotherapies when targeting CD70 for AML.

We additionally gained important insights into optimizing bicistronic CAR design through this work. We show that the greatest *in vitro* and *in vivo* efficacy was seen in constructs that contained both a 4-1BB and CD28 costimulatory domains on separate CARs, which may suggest an important finding for future bicistronic CAR designs. Indeed, a significant body of literature has shown that CARs utilizing CD28 or 4-1BB costimulatory domains activate different, but complementary, signaling pathways^74,75^, and that combining both signaling domains in the same cell can improve potency and antitumor efficacy, potentially by combining the early proliferation and activation of the CD28 domain with the prolonged persistence of the 4-1BB domain^76–78^. It appears our bicistronic CAR design is also able to take advantage of this synergistic signaling, as designs BCD5 and BCD6, which contained both costimulatory domains, displayed both earlier expansion and longer persistence when compared to BCD1, which contained only the 4-1BB domain on both CARs expressed.

Furthermore, we note that the previously documented exceptionally high *in vivo* CAR-T expansion phenotype, observed by several groups utilizing the CD27 natural ligand-based anti- CD70 CAR^59,67,68^, is maintained in this bicistronic CAR format. The relative strength of this expansion appears markedly stronger with a CD28 costimulatory domain coupled to the CD27- based natural ligand CAR, as seen in the BCD5 and BCD6 designs, compared to a 4-1BB costimulatory domain, as in BCD1. Although the exact mechanism of this enhanced *in vivo* expansion is not yet well understood, prior work has shown that only the extracellular CD27- based binder is necessary for this phenotype, as swapping the transmembrane or signaling domains does not ablate the proliferative advantage. This finding also suggests that combining the CD27-based CAR with other CARs, either for AML or other tumor contexts, could serve as a strategy to boost the proliferation and persistence of CAR-T products that show poor persistence as a monotherapy.

We also acknowledge that this study contains several limitations. Notably, we characterize co-expression of CD70 and aITGB2 on 15 primary AML samples, which is sufficient to see both a significant effect size and statistical significance in support of a dual-antigen targeting approach. However, this sample size is insufficient to characterize differences in expression between different AML subtypes, and future work will analyze a larger number of AML samples to further explore these relationships. Furthermore, for several of the patient samples analyzed, there appears to be a small population of CD70^-^ aITGB2^-^ AML blasts, suggesting that our specific bicistronic CAR design may not eliminate all blasts in those patients and potentially allow for antigen-negative relapse. However, we also note that identification of AML blasts by flow cytometry-based surface expression is not perfect, and that some of these cells may actually represent a healthy progenitor cell population, or that blasts which do not have detectable antigen expression signals by flow cytometry may still express some degree of target antigen and could still be therapeutically targeted. Additionally, significant past literature has shown that the hinge and transmembrane regions of CARs, as well as alternative T-cell signaling architecture, can play a significant role in their cytotoxic ability^79–83^. In this study, each CAR was only tested in a single hinge and transmembrane framework, but future work will focus on evaluating the optimal CAR design for non-signaling regions of the CARs as well.

Overall, this work identifies a novel dual antigen-targeting therapy against the AML-specific antigens CD70 and aITGB2 which is shown to overcome current concerns with target antigen heterogeneity and off-tumor toxicity in anti-AML CARs. These studies strongly support further preclinical development of this therapy, with the potential for future translation to the clinic.

## Methods

### CD70 and ITGB2 Expression Analysis

Data was analyzed from previously published bulk RNA-seq datasets (TCGA AML, BEAT AML and TARGET AML), available under accession codes phs000178 (https://www.ncbi.nlm.nih.gov/projects/gap/cgi-bin/study.cgi?study_id=phs000178), phs001657 (http://www.ncbi.nlm.nih.gov/projects/gap/cgi-bin/study.cgi?study_id=phs001657), and phs000218 (http://www.ncbi.nlm.nih.gov/projects/gap/cgi-bin/study.cgi?study_id=phs000218), respectively, through the NIH/NCI GDC Data Portal.

### Cell Lines, PDX, and Human Samples

NOMO-1 and MOLM-14 cell lines were obtained from DSMZ and THP-1 cell lines were obtained from ATCC. All cell lines were grown in RPMI-1640 medium (Gibco, 11875093) supplemented with 20% fetal bovine serum (FBS; BenchMark, Gemini, Lot # 100-106) and 100 U/mL penicillin/streptomycin (UCSF Cell Culture Facility). All cells were grown at 37 °C with 5% CO2. All AML PDXs were procured from the PRoXe repository^63^ at Dana–Farber Cancer Institute under an appropriate materials transfer agreement. Primary AML samples were obtained from the UCSF Hematologic Malignancies Tissue Bank and the Pediatric Hematopoietic Tissue Cell Bank.

### Generation of *ITGB2* and *CD70* knockout cells

Knockout cell lines or primary cells were generated using *in vitro* nucleofection of Cas9 ribonuclease protein complexed to sgRNA against the gene of interest. Unless otherwise specified, 2 μl of each sgRNA (100 µM; Synthego Corporation) and recombinant Cas9 protein (40 µM; QB3 MacroLab, University of California, Berkeley) was incubated at 37 °C for 15 min to generate ribonuclease complex, which was then nucleofected using a 4D-Nucleofector (Lonza) with the built-in program DS-137 for cell lines (using Lonza V4XC-2032) unless otherwise specified and EO-115 for primary T cells (using Lonza V4XP-3032). Knockout cell lines were purified through Fluorescent Activated Cell Sorting (FACS) on a BD FACS Aria Fusion instrument.

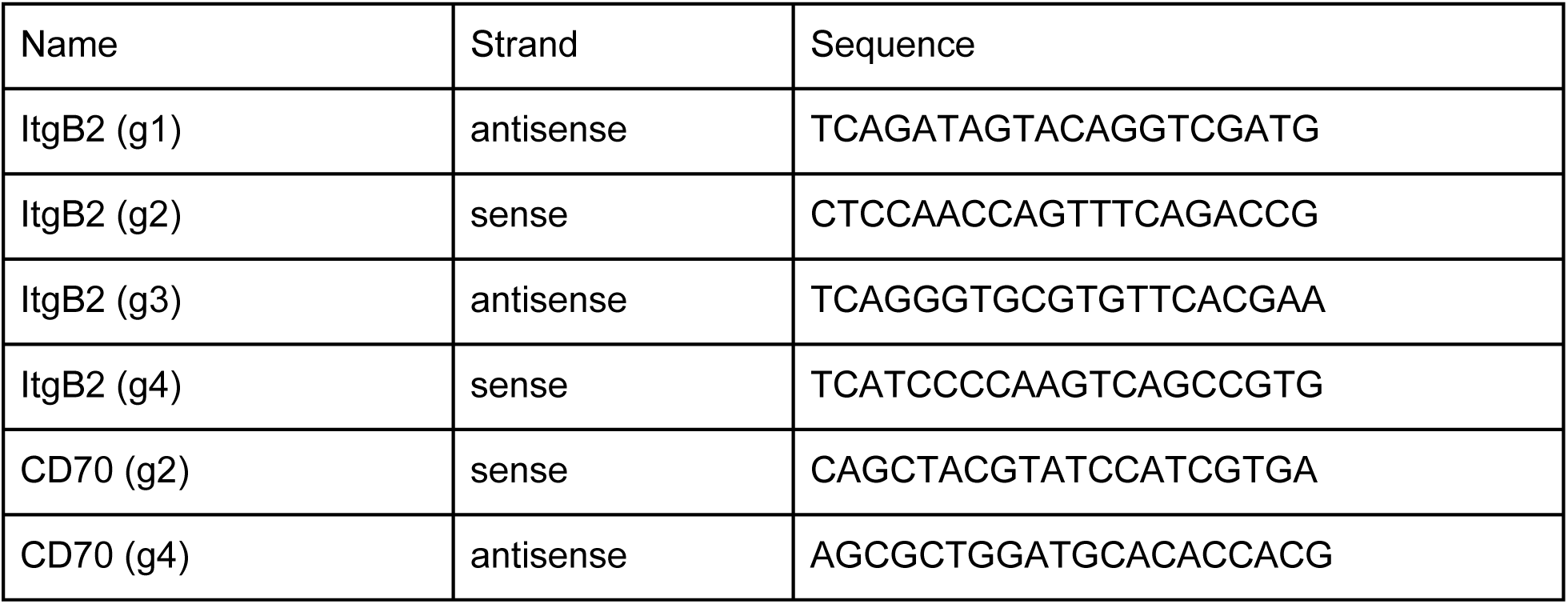

### Genomic DNA confirmation of CRISPR-Cas9 knockouts

DNA extraction from 1 million cells, whether fresh or frozen, was conducted using the Monarch® Genomic DNA Purification Kit (NEB, Cat # T3010S) or Zymo DNA Clean & Concentrator-5 (Zymo Research, D4013). Primers were procured from Integrated DNA Technologies to yield a ∼500 bp amplicon, ensuring a minimum of 200 bp upstream and downstream from the target sgRNA sequence. The primers were reconstituted in nuclease-free water to create a 10 µM stock. A PCR reaction employing 0.5 µM of both forward and reverse primers was carried out using the NEBNext® High-Fidelity 2X PCR Master Mix (Cat # M0541S) with 1 ng to 1 µg of genomic DNA template. Subsequently, the PCR product was purified using the QIAquick PCR Purification Kit (QIAGEN, Cat # 28104). Sanger Sequencing was performed by Quintara Biosciences. The resulting AB1 files for both control and experimental DNA groups were uploaded to the Synthego ICE analysis tool (https://ice.synthego.com/), where indel percentage, Model Fit (R^2^), and Knockout Score were employed to assess knockout efficiency.

### Molecular Cloning and DNA Plasmids

Genes encoding the CAR T antigen binding sequences were synthesized as gene fragments from Twist Biosciences (South San Francisco, CA). DNA fragments were then cloned into a lentiviral expression vector via NEB HiFi assembly (NEB, E2611S). Whole plasmid DNA sequencing was performed to confirm accuracy of the vector. This final construct was then expressed in Stbl3 Competent E. coli (Macro Lab, UC Berkeley, CA). DNA was isolated using either QIAGEN Plasmid Plus Midi Kit or QiaPrep Spin Miniprep Kit (Qiagen, 27104).

### Flow Cytometry

Immunostaining of cells was performed as per the instructions from the antibody vendor unless stated otherwise. One million cells were resuspended in 100 µl of FACS buffer (PBS + 2% FBS) with 5µL of human Fc Block (Biolegend, 422302) added. Cells were incubated at room temperature for 10 min, then 1-5µL of antibody was added (based on prior titration). Cells were incubated at 4°C for 30 minutes and washed three times with FACS buffer before flow cytometry analysis. For staining the active form of integrin β2, the antibody incubation step was performed at 37 °C for 1 hour, before proceeding with the regular antibody staining. For staining primary AML cells for activated integrin β2, the FACS buffer was RPMI-1640 + 5% FBS + 2% bovine serum albumin (BSA) + 50 µg/ml DNase I (Gold Biotechnology, D-301-500). Compensation used UltraComp eBeads Compensation Beads (Invitrogen, 01-2222-42). Flow cytometry analysis was performed on the CytoFLEX platform (Beckman Coulter), and data were analyzed using FlowJo_v10.10.0.

### Antibody Inventory

| Company | Catalog # | Clone | Antibody |
| --- | --- | --- | --- |
| Biolegend | 355106 | 113-16 | FITC anti-human CD70 |
| Biolegend | 355122 | 113-16 | APC/Fire750 anti-human CD70 |
| Biolegend | 355104 | 113-16 | PE anti-human CD70 |
| Biolegend | 355110 | 113-16 | APC anti-human CD70 |
| Biolegend | 363408 | m24 | Brilliant Violet 421 anti-human CD11a/CD18 (LFA-1) (ItgB2) |
| Biolegend | 303404 | WM53 | PE anti-human CD33 |
| Biolegend | 303422 | WM53 | Brilliant Violet anti-human CD33 |
| Biolegend | 303304 | HIM3-4 | FITC anti-human CD33 |
| Biolegend | 343504 | 581 | FITC anti-human CD34 |
| Biolegend | 343535 | 581 | APC/Fire750 anti-human CD34 |
| Biolegend | 368512 | 2D1 | APC anti-human CD45 |
| Biolegend | 356620 | HB-7 | Brilliant Violet 650 anti-human CD38 |
| Biolegend | 303506 | HIT2 | PE anti-human CD38 |
| Biolegend | 329916 | EH12.2H7 | Pacific Blue anti-human CD279 (PD-1) |
| Biolegend | 329705 | 29E2A3 | PE anti-human CD279 (PD-1) |
| Invitrogen | 17-2239-41 | 3DS223H | APC anti-human CD223 (LAG-3) |
| Invitrogen | 17-3109-42 | F38-2E2 | APC anti-human CD366 (TIM-3) |
| Invitrogen | 12-0458-42 | H100 | PE Anti-Hu CD45RA |
| Biolegend | 344824 | SK7 | Pacific Blue anti-human CD3 |
| Biolegend | 982416 | HIB19 | APC/Fire750 anti-human CD19 |
| Biolegend | 982508 | M5E2 | PE anti-human CD14 |
| Biolegend | 367166 | 63D3 | FITC anti-human CD14 |
| Biolegend | 306708 | IP26 | PE anti-human TCR a/b |
| Biolegend | 637310 | L5 | PE anti-DYKDDDDK (flag) Tag |
| Biolegend | 637322 | L5 | Brilliant Violet 421 anti-DYKDDDDK (flag) Tag |
| Cell Signaling Technologies | 2233S | 9B11 | Myc-Tag Mouse mAB (Alexa ® 647 Conjugate) |
| Biolegend | 422302 | N/A | Human TruStain FcX |

### Human primary T cell isolation

Primary T cells were isolated from LeukoPaks (Stem Cell Technologies, 200-0092) using an EasySep human T cell isolation kit (StemCell, 17951) based on magnetic bead separation.

### Human Primary Acute Myeloid Leukemia (AML) Sample Processing

Primary AML samples were sourced from the Hematologic Malignancies Tissue Bank at UCSF. Patient samples were collected under University of California, San Francisco (UCSF) Institutional Review Board-approved protocols. Samples were spun down at 500✕g for 5 minutes and resuspended in 1X Ammonium Chloride (ACK) lysis buffer, then gently mixed on a rocker at room temperature for 10 minutes. Samples were then spun at 500✕g for 5 minutes and then resuspended in 5mL of Bulk Lyse Wash Solution (PBS + 0.1% Sodium Azide + 0.5% Bovine Serum Album (BSA)) and run through a 70µm filter. Samples were centrifuged at 500✕g for 5 minutes and resuspended in FACS Buffer (PBS + 2%(v/v) FBS) and cell counts were taken before proceeding with antibody staining.

### PDX and Primary AML Co-Culture Assay

HS-5 bone marrow stromal cells (ATCC HS-5, CRL-3611) were grown in RPMI-1640 medium (Gibco, 11875093) + 10% FBS + 100 U/ml penicillin–streptomycin. HS-5 were seeded at 50% confluency 2 days prior to starting the co-culture assay. Confluency was confirmed to be 70- 80% via brightfield microscope immediately prior to co-culture. Frozen Primary or PDX samples were thawed in 20% RPMI-1640 medium (Gibco, 11875093) + 20% FBS + 100 U/ml penicillin– streptomycin + HEPES + GlutaMAX (Gibco™. Cat # 35050061) + 10 µg/ml DNase I and spun at 200✕g for 5 minutes. Samples were resuspended in 20% FBS RPMI-1640 medium (20% FBS + 100 U ml–1 penicillin–streptomycin + HEPES + GlutaMAX (Gibco™. Cat # 35050061). The primary AML or PDX was added directly onto the 70-80% confluent HS-5s at a concentration of 1 million cells/mL. Cytokines interleukin 3 (IL-3), stem cell factor (SCF), and FMS-like tyrosine kinase 3 ligand (FLT3L) were added at a final concentration of 25 ng/mL each. Control co-cultures were grown without HS-5s or without cytokines. Primary AML or PDX were transferred to fresh HS-5 with cytokines every two days and cultured at 1 million cells/mL. Flow cytometry was performed on 500,000 cells per sample immediately after thaw as well as at multiple timepoints thereafter, up to 7 days after thaw.

### Lentiviral Vector Production

Lenti-X 293T cells (Takara, Cat # 632180) were transfected with CAR expression plasmid and Mirus Bio™ TransIT™-Lenti Transfection Reagent (Cat # MIR6600) or Invitrogen™ Lipofectamine™ 3000 Transfection Reagent (Cat # L3000015) and ViralBoost Reagent (Alstem Cell Advancements, Richmond, CA. Cat # VB100). Lenti-X 293T cells were cultured for 2-3 days, and then lentivirus was harvested at 24h and 48h and concentrated according to the Lenti-X Concentrator (Takara Bio, 631232) protocol.

### CAR T-cell Production and Expansion

CD3-isolated T-cell populations were thawed and seeded in CTS OpTmizer medium with CTS supplement (Thermo Scientific, A1048501) supplemented with 5% human AB serum (HP1022; Valley Medical), 1% GlutaMAX (Gibco™. Cat # 35050061), and 100 U/mL penicillin/streptomycin (Fisher Scientific, 15-140-122) at a concentration of 1 million cells/mL. Following overnight recovery, T-cells were nucleotides with Cas9-RNP complex with sgRNA targeting the *CD70* and *ITGB2* genes as per the nucleofection protocol above. Following overnight recovery after CRISPR editing, T cells were stimulated with CD3/CD28 Dynabeads (11131-D; Thermo Fisher Scientific) according to the manufacturer’s instructions (25 μL of beads per 1 million T-cells) for 4 days and grown in the presence of recombinant interleukin-7 (PeproTech, 200-07) and interleukin-15 (PeproTech, 200-15) at 10 ng/mL. Transduction with CAR lentivirus was performed 1 day after the start of bead stimulation, using approximately 100 microliters of concentrated lentivirus per million T-cells. After the removal of CD3/CD28 activation beads, transduction efficiency was assessed by flow cytometry. CAR-T cells were labeled with intracellular GFP or myc/flag tags to allow assessment of CAR transduction efficiency.

### *In Vitro* Cytotoxicity Assay

AML cell lines were engineered to stably express luciferase using lentiviral transduction. For each cell line, 50,000 target tumor cells were seeded in 50 uL of RPMI-1640 media in a white 96 well flat-bottom plate (Greiner Bio-One). Subsequently, an appropriate number of CAR-T cells were added at various effector:target ratios in 50uL of T-cell media with *n*=3 replicates per condition. This coculture was incubated at 37 C for 18-24 hours. The next day, 100uL of D-Luciferin (Gold Biotechnology, LUCK-1G) was added to each well to a final concentration of 375 μg/ml, followed by luminescence detection using GloMax Explorer (Promega). The bioluminescence readings were averaged amongst the *n*=3 technical replicates for each construct and ratio. Bioluminescence readings were normalized to the negative control condition for each E:T ratio and cell line to create a cytotoxicity scale of 0-100.

#### Murine AML Cell Line CAR-T Studies

Murine studies were conducted in accordance with UCSF Institutional Animal Care and Use Committee protocol and guidelines. NSG (NOD.Cg-*Prkdc^scid^ Il2rg^tm1Wjl^*/SzJ, strain #:005557, Jackson Laboratories) male mice, 6-8 weeks old, were bred at the UCSF Breeding Core Facility. A mixed population of 500,000 *CD70* KO THP-1 and 500,000 *ItgB2* KO luciferase-expressing THP-1 cells were injected intravenously via the tail vein. Three days following tumor injection, mice were transfused intravenously with 1.5 million of CAR-T cells or untransduced T cells. Bioluminescence imaging and peripheral blood draws were done weekly to assess tumor burden (Perkin Elmer In Vivo Imaging System, Caliper Life Sciences). Flow cytometry was performed on the blood draws to monitor CAR-T proliferation and T-cell phenotypes. Survival end point of the study was determined by signs of symptomatic illness in the animals and euthanasia humane endpoints.

#### Murine AML PDX Model CAR-T Studies

Murine studies were conducted in accordance with UCSF Institutional Animal Care and Use Committee protocol and guidelines. NOG-EXL (NOD.Cg-*Prkdc^scid^ Il2rg^tm1Sug^* Tg(SV40/HTLV- IL3,CSF2) 10-7Jic/JicTac, model no. HSCCB-13395, Taconic Biosciences) female mice, 6-8 weeks of age, were purchased from Taconic Biosciences for this study. Mice were preconditioned with 2 doses of busulfan, each at 12.5 mg/kg, for a total dose of 25 mg/kg. Two days later, 2 million AML PDX cells were resuspended in PBS and injected via tail vein. Ten days later, 1.5 million CAR T cells were injected IV via tail vein. Peripheral blood draws were performed weekly, with flow cytometry analysis used as readouts for tumor burden and CAR-T proliferation. Survival end point of the study was determined by signs of symptomatic illness in the animals and euthanasia humane endpoints. Spleens were harvested according to primary sample processing protocols.

#### Colony Forming Assay

CD34+ cells (1 × 10^3^) were isolated from healthy donor GM-CSF-mobilized peripheral blood (Stemcell Technologies EasySep Human CD34 Positive Selection Kit II) and co-incubated with CAR-T cells, untransduced T-cells, or medium only (IMDM, 2% FBS and penicillin/streptomycin) at a 5:1 E:T ratio for 5 h in V-bottom 96-well plates in triplicate. Cells were then transferred to a total volume of 1.1 mL of methylcellulose-based medium (MethoCult H4434 Classic, StemCell Technologies) per condition in a 35 mm dish and placed in a 37°C cell culture incubator for 14 days, maintaining 100% humidity to prevent the media from drying out. Colonies were counted and classified as granulocyte-erythroid-macrophage-megakaryocyte colony forming unit (CFU-GEMM), granulocyte-macrophage colony forming unit (CFU-GM), burst-forming unit-erythroid (BFU-E), or erythroid colony forming unit (CFU-E). Images were acquired with a Keyence microscope using ×10, brightfield and color mode and averaged from triplicate conditions.

#### CD34+ Cytotoxicity Assay

CD34+ cells previously isolated from healthy donor GM-CSF mobilized blood were thawed and plated at 25,000 cells per well in IMDM + 2% FBS media and co-incubated with CAR-T cells or untransduced T-cells at a 3:1 E:T ratio in T-cell media in a final volume of 200 μL per well. Each condition was performed in triplicate. After 24 hours of incubation at 37°C, co-cultures were transferred to a 96 well round bottom plate and stained with a fixable viability dye and antibodies for CD34 and CD3, along with counting beads to allow for accurate quantification of live CD34+ cells by flow cytometry. The number of live CD34+ cells was normalized to a negative control condition with no T-cells added and a positive control condition with 1% Tween-20 added to calculate percent cytotoxicity on a 0-100 scale, averaged between replicates for each condition.

### Study Design

Study design has generally been described for each experiment in the results section. Minimum sample sizes for murine studies were selected based on power calculations for improved survival based on an α of 0.05, a β of 0.2, and an effect size of 7 days of improved survival. All data from experiments with appropriately behaving positive and negative controls were included, and no outliers were excluded in our analyses. Most experiments were performed in at least 3 experimental replicates, and biological replicates, such as multiple T-cell donors and AML cell line models, were used throughout the study.

### Statistical Analyses

All hypotheses were tested at an α=0.05 level unless otherwise indicated. Data was analyzed using GraphPad Prism 10, and specific statistical tests used for each experiment have been indicated within the main text or figure legends.

## Acknowledgements

We thank blood donors, patients, and their families for participating in research and contributing primary specimens for research purposes

## Author Contributions

Conceptualization: ASK, HJ, APW

Experimentation and Analysis: ASK, HJ, AI, CK, NC, JR, PP, JACS, FS, VS, BJH

Patient Sample Acquisition: NL, JW, NR, ACL

Supervision: JE, APW

Manuscript Writing: ASK, HJ, JE, APW

Figure Design: ASK, HJ, BJH

## Data and Materials Availability

Bulk RNA-seq datasets (TCGA AML, BEAT AML and TARGET AML) used for patient target antigen analysis is available under accession codes phs000178 (https://www.ncbi.nlm.nih.gov/projects/gap/cgi-bin/study.cgi?study_id=phs000178), phs001657 (http://www.ncbi.nlm.nih.gov/projects/gap/cgi-bin/study.cgi?study_id=phs001657), and phs000218 (http://www.ncbi.nlm.nih.gov/projects/gap/cgi-bin/study.cgi?study_id=phs000218), respectively, through the NIH/NCI GDC Data Portal. All other data are available in the main text or the supplementary materials

---

**Supplementary Figure 1.**
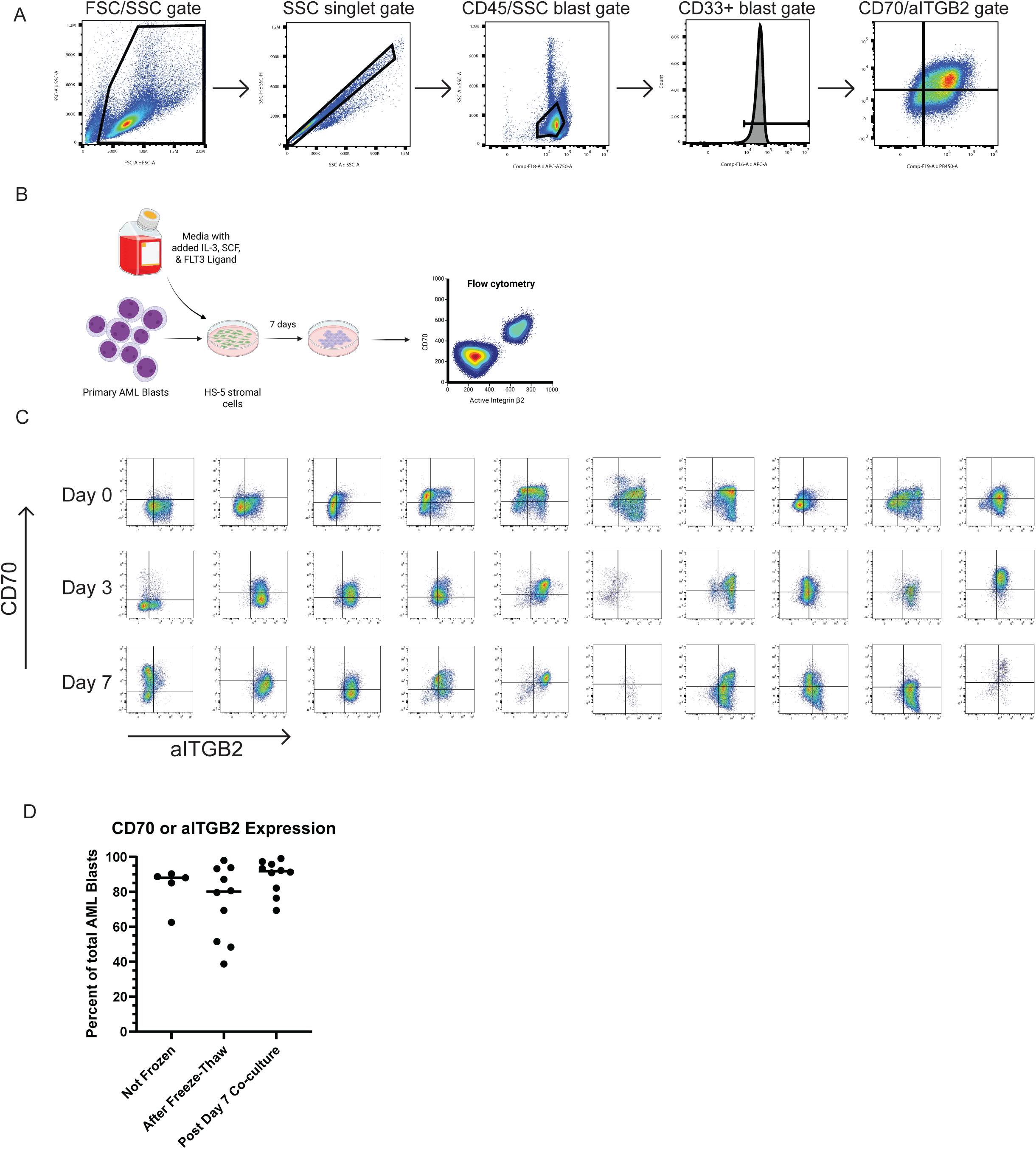
Additional AML primary sample flow cytometry strategy and data: **A)** Representative gating strategy for CD70 and aITGB2 expression on AML blasts isolated from primary sample peripheral blood or bone marrow. B) Cartoon diagram of *ex vivo* AML primary sample co-culture system. C) Flow cytometry plots of CD70 and aITGB2 expression for N = 10 frozen AML primary sample immediately upon thaw (top) and following 3 (middle) and 7 (bottom) days of co-culture with HS-5 bone marrow stromal cell line and supportive cytokines. D) Percent of AML blasts captured in a CD70/aITGB2 OR-targeting gate compared between AML samples that were never frozen, immediately after a freeze-thaw event, or after 7 days of co-culture with stromal cell line and cytokines.

**Supplementary Figure 2.**
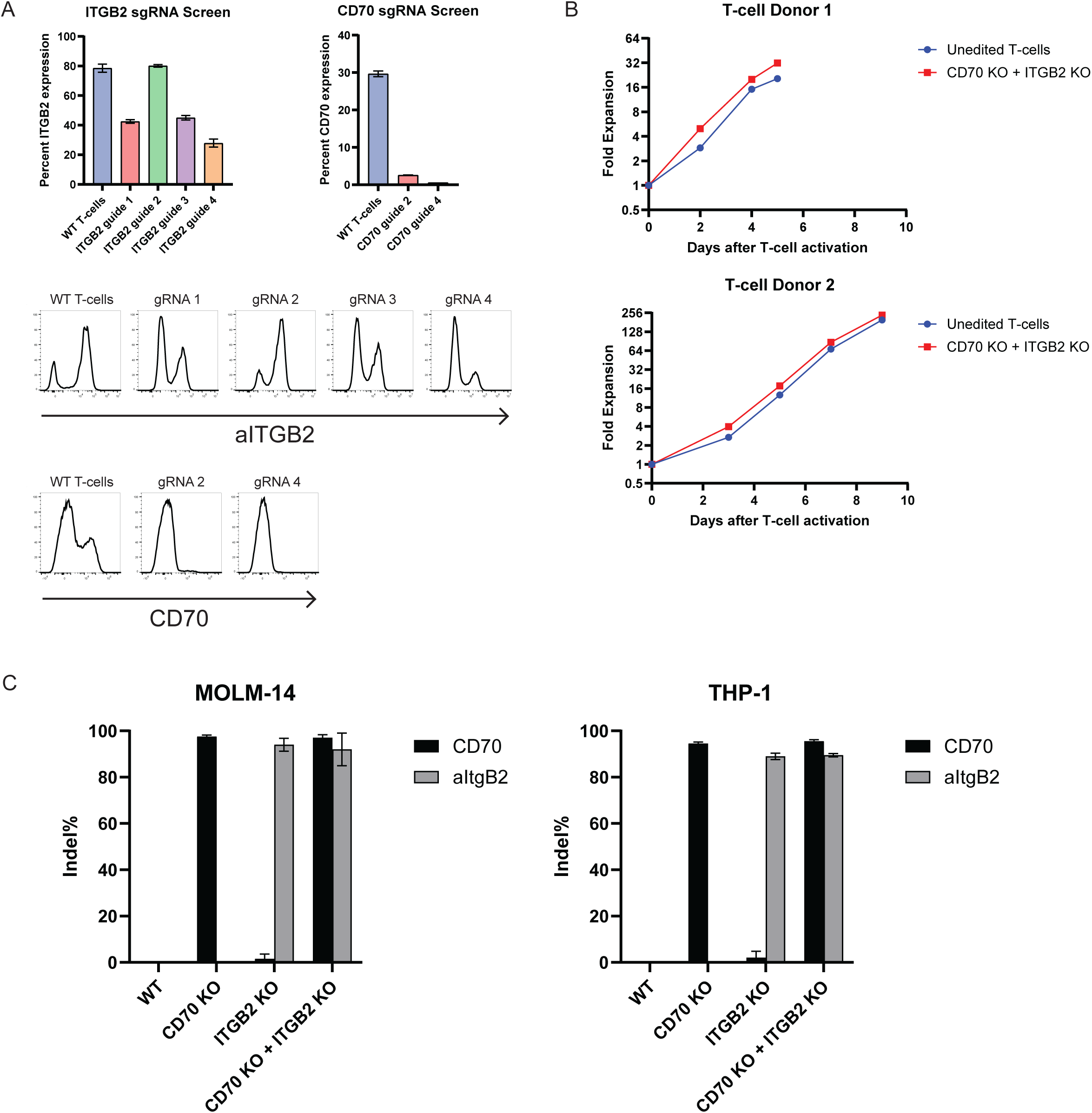
CD70 and ITGB2 knockout is effective and well-tolerated in T-cells: **A)** Top-Remaining percent of T-cells expressing aITGB2 or CD70 four days after electroporation with a complex of Cas9 and a gRNA targeting the indicated gene. Bottom-representative flow cytometry histograms of T-cell antigen expression for each gRNA tested. B) Relative expansion of CD3/CD28 bead-activated T-cells following CRISPR knockout of CD70 and aITGB2 compared to unedited T-cells. C) Percent of each cell line confirmed to have a genetic knockout at each target gene locus following genomic DNA sequencing, as measured by the Synthego ICE tool. Error bars represent SEM.

**Supplementary Figure 3.**
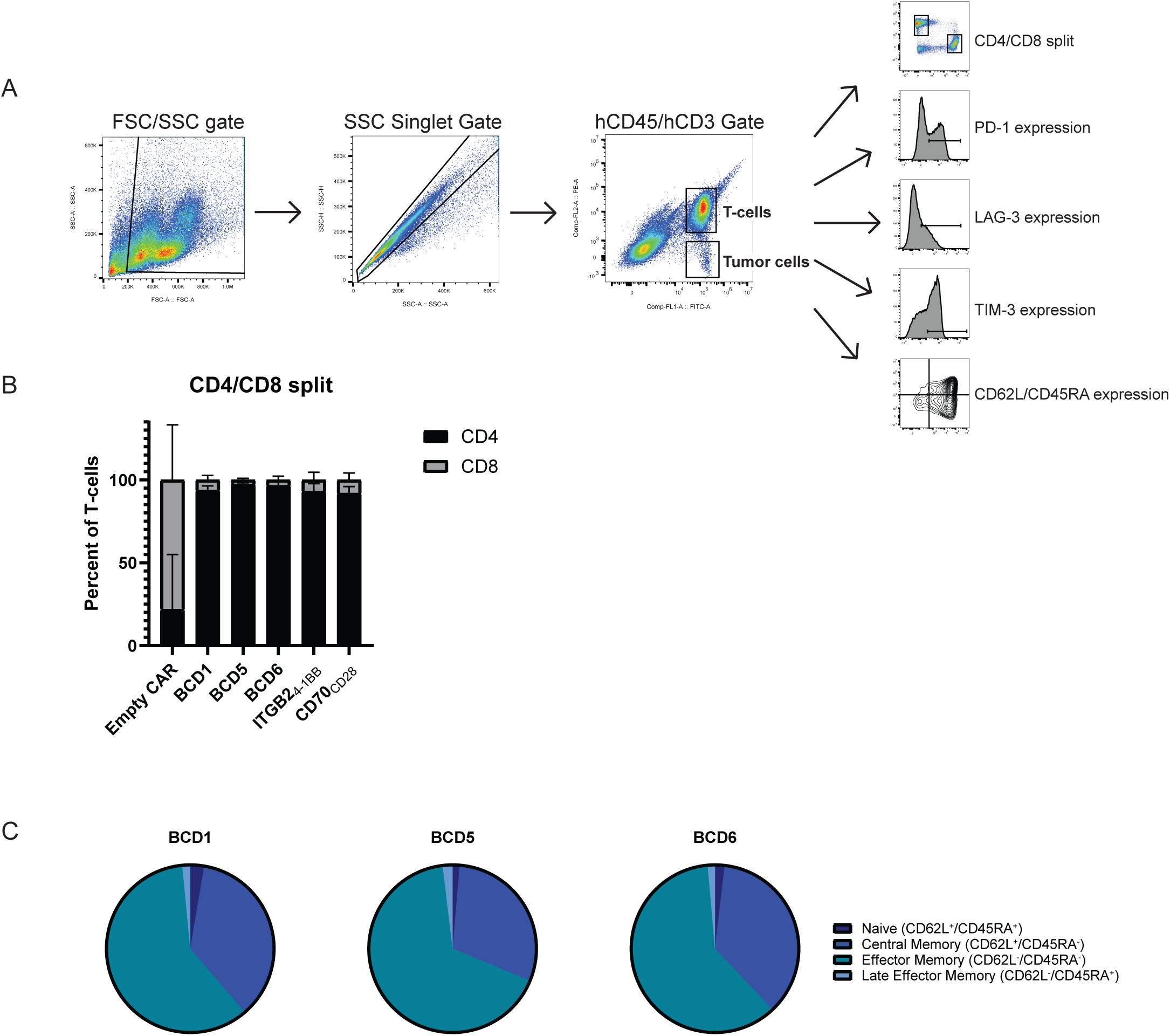
Additional flow cytometry strategy and data from *in vivo* antigen heterogeneity experiment: **A)** Representative gating strategy for flow cytometry on weekly mouse peripheral blood draw samples to evaluate T-cell and tumor cell phenotypes and surface protein expression. **B)** Percent of CD3^+^ T-cells in the peripheral blood expressing CD4 or CD8, 34 days after CAR-T injection. Data collected from N = 3 mice per group by flow cytometry, error bars represent SD. **C)** Relative proportion of CAR-Ts classified into each memory/effector T-cell phenotype, 55 days after CAR-T infusion into mice. Classification assessed by flow cytometry of CD62L and CD45RA expression on CAR-T cells isolated from peripheral blood. Data presented is the average of N = 3 mice evaluated per treatment group.

**Supplementary Figure 4.**
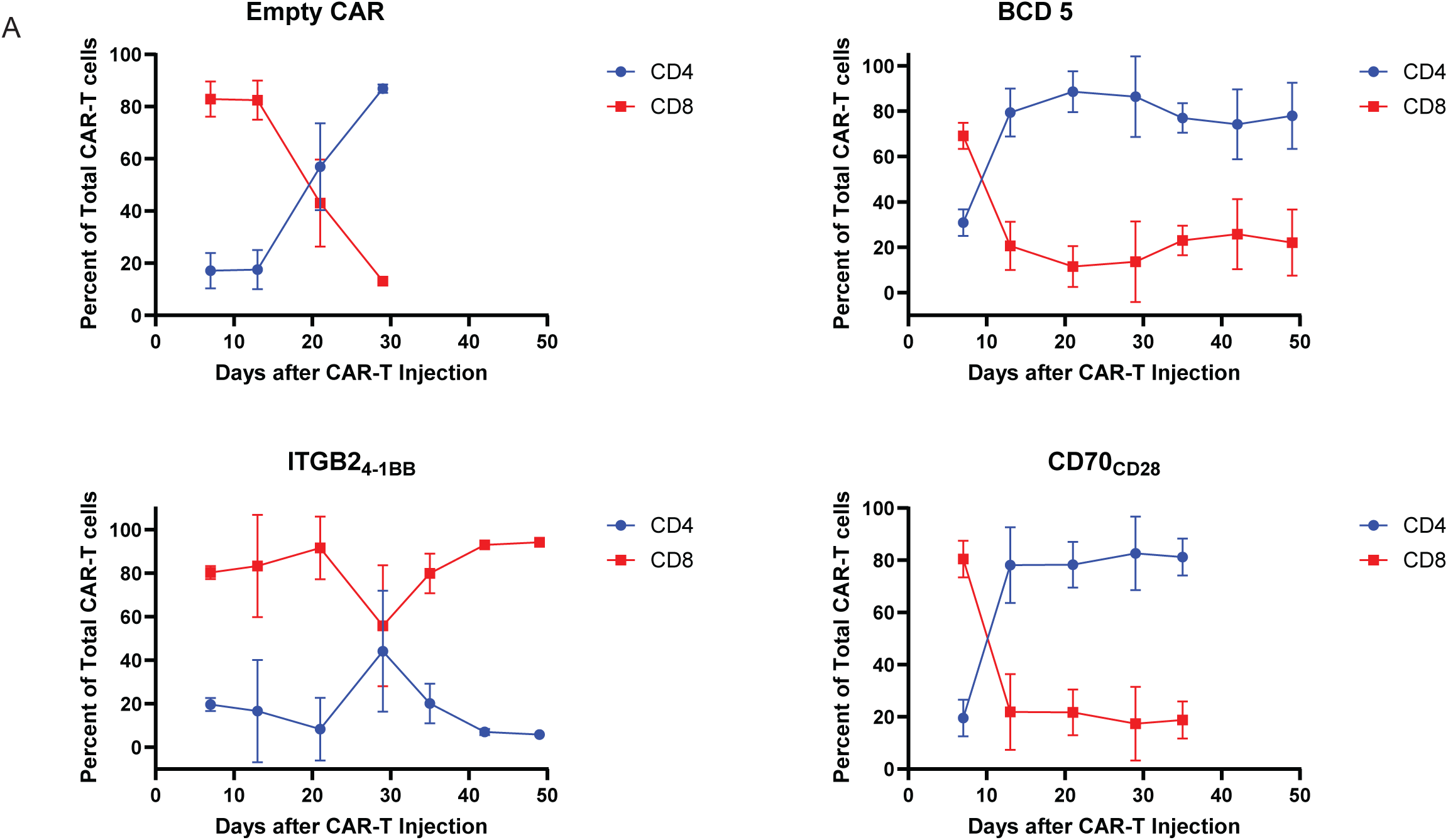
Long-lived CAR-Ts for BCD5 are primarily CD4+: **A)** Percent of CD3^+^ T-cells in the peripheral blood expressing CD4 or CD8 over time. Data collected from N = 3 mice per group at each time point by flow cytometry, error bars represent SD.

